# Rapid and site-specific deep phosphoproteome profiling by data-independent acquisition (DIA) without the need for spectral libraries

**DOI:** 10.1101/657858

**Authors:** Dorte B. Bekker-Jensen, Oliver M. Bernhardt, Alexander Hogrebe, Ana Martinez del Val, Lynn Verbeke, Tejas Gandhi, Christian D. Kelstrup, Lukas Reiter, Jesper V. Olsen

## Abstract

Quantitative phosphoproteomics has in recent years revolutionized understanding of cell signaling, but it remains a challenge to scale the technology for high-throughput analyses. Here we present a rapid and reproducible phosphoproteomics approach to systematically analyze hundreds of samples by fast liquid chromatography tandem mass spectrometry using data independent acquisition (DIA). To overcome the inherent issue of positional phosphopeptide isomers in DIA-based phosphoproteomics, we developed and employed an accurate site localization scoring algorithm, which is incorporated into the Spectronaut software tool. Using a library of synthetic phosphopeptides spiked-in to a yeast phosphoproteome in different ratios we show that it is on par with the top site localization score for data-dependent acquisition (DDA) based phosphoproteomics. Single-shot DIA-based phosphoproteomics achieved an order of magnitude broader dynamic range, higher reproducibility of identification and improved sensitivity and accuracy of quantification compared to state-of-the-art DDA-based phosphoproteomics. Importantly, direct DIA without the need of spectral libraries performed almost on par with analyses using specific project-specific libraries. Moreover, we implemented and benchmarked an algorithm for globally determining phosphorylation site stoichiometry in DIA. Finally, we demonstrate the scalability of the DIA approach by systematically analyzing the effects of thirty different kinase inhibitors in context of epidermal growth factor (EGF) signaling showing that a large proportion of EGF-dependent phospho-regulation is mediated by a specific set of protein kinases.

## INTRODUCTION (Brief and focused)

Site-specific protein phosphorylation is one of the most important post-translational modifications (PTMs) as it can rapidly modulate a protein’s function by changing its activity, subcellular localization, interactions or stability^1^. It is a highly dynamic modification that regulates essentially all cellular signaling networks. Deregulated phospho-signaling is therefore a hallmark of cancer and many other diseases. Major advances in phosphopeptide enrichment strategies, instrument performance and computational analysis tools have made mass spectrometry-based phosphoproteomics the method of choice for the study of protein phosphorylation on a global scale. Large-scale quantitative phosphoproteomics has proven to be successful in addressing unsolved questions in cell signaling and biomedicine^2–5^. However, the majority of successful phosphoproteomics studies typically involves days or even weeks of measurements by liquid chromatography tandem mass spectrometry (LC-MS/MS) to analyze few cellular conditions with sufficient depth to identify and pinpoint the functional phosphorylation sites. Moreover, current tandem mass spectrometric sequencing speed and the semi-stochastic nature of data-dependent acquisition (DDA) make it challenging for phosphoproteomics to systematically and reproducibly analyze phosphorylation sites across large numbers of samples. This limits its application for high-throughput applications such as drug screening.

With the advent of fast scanning high-resolution tandem mass spectrometers, DIA has appeared as powerful alternative to DDA in shotgun proteomics^6–8^. In a DIA analysis, all (phospho)peptides within a predefined mass-to-charge (m/z) window are co-fragmented and the resulting fragments measured together. This analysis is repeated as the mass spectrometer goes through the full mass range, which facilitates systematic measurement of all peptide ions regardless of their intensity and overcomes the precursor selection problem of DDA. DIA typically provides broader dynamic range, higher peptide identification rates, improved reproducibility of identification, and accuracy for quantification. However, due to the multiplexed fragment ion spectra, DIA requires more elaborate data processing algorithms and software solutions for spectral deconvolution, which typically make use of pre-recorded spectral libraries. Moreover, apart from the unambiguous identification of the phosphopeptide sequence, the deconvoluted tandem mass spectra should contain sufficient information to localize phosphorylation sites with single amino acid resolution^1^.

To address these issues, we have developed an optimized label-free quantitative phosphoproteomics approach combining fast liquid chromatography tandem mass spectrometry with site-specific data-independent acquisition (DIA). This approach allowed us to systematically and reproducibly analyze more than ten thousand phosphorylation sites across hundreds of samples. We developed and employed algorithms to accurately localize phosphorylation sites in DIA datasets and determine their fractional stoichiometry on a system-wide scale. We apply this strategy to identify phosphorylation site targets of ten major protein kinases in the epidermal growth factor signaling pathway^9^.

## RESULTS (General description of the method followed by its validation)

### Comparison of DDA and DIA for large-scale quantitative phosphopeptide analysis

To enable large-scale phosphoproteomics studies with increased depth and throughput, it is necessary to reduce the amount of input protein, improve workflow reproducibility and decrease mass spectrometry instrument time usage. With this in mind we optimized a fast and scalable single-shot analysis workflow based on high-throughput magnetic Ti-IMAC bead enrichment of phosphopeptides from 200 ug of starting tryptic peptide material (**Figure 1A**). With this approach we routinely quantify ~7000 phosphopeptides in just 15 min of LC-MS/MS analysis time with fast 28 Hz Higher-energy Collisional Dissociation (HCD)^10^ scanning method on a Q Exactive HF-X mass spectrometer^8^ with overall MS/MS identification rates of more than 50%. This is close to 500 unique phosphopeptides per minute of gradient time, which is significantly more identifications than the commonly-used TiO2-based workflow^8^.

**Figure 1.**
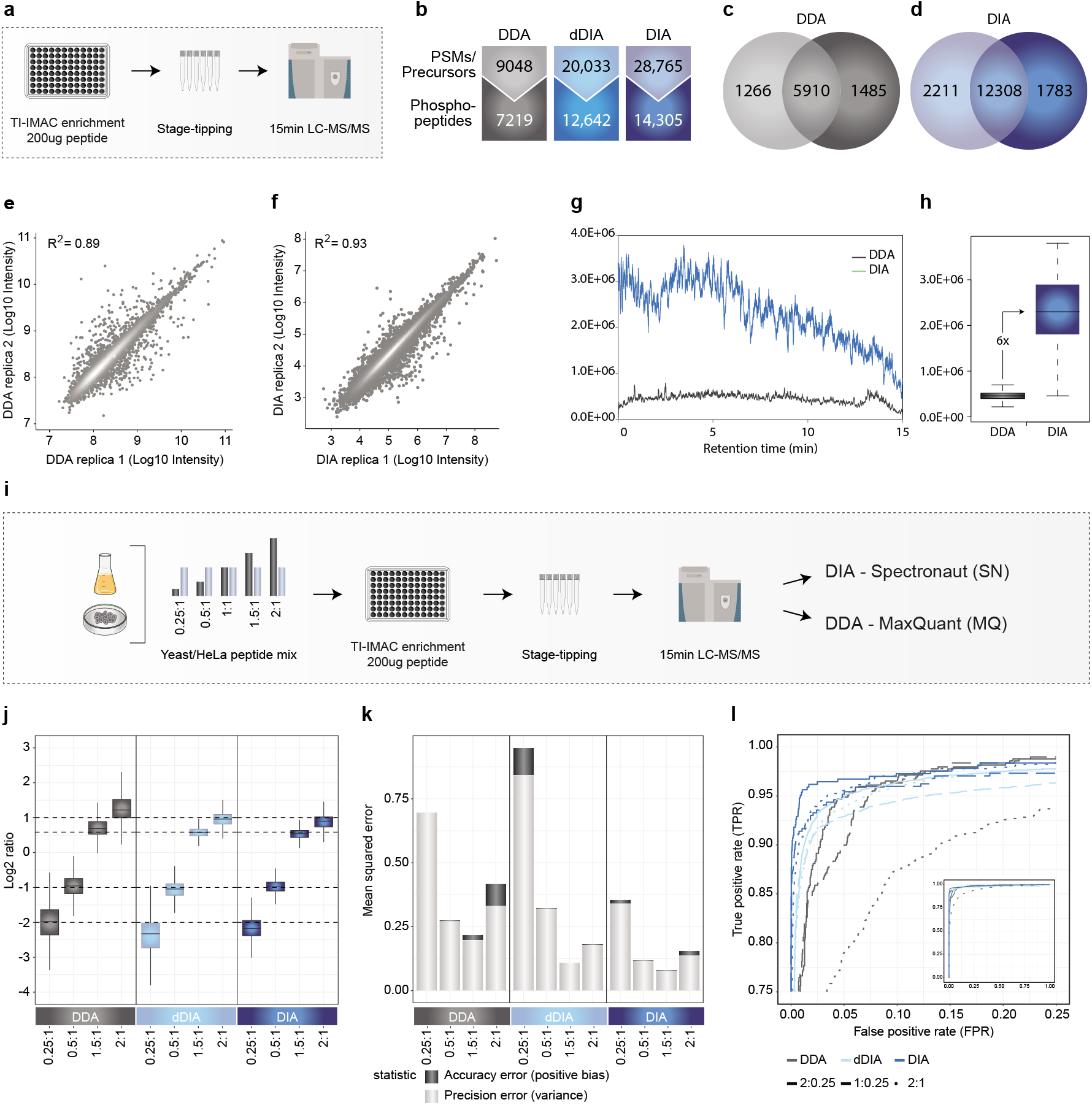
High-throughput and sensitive phosphoproteomics for DDA & DIA – identification and quantification **(a)** Experimental workflow for phosphoproteomics. **(b)** Comparison of quantified phosphopeptides with DDA & DIA. **(c)** Overlap of phosphopeptides between two replica with DDA. (d) Overlap of phosphopeptides between two replica with DIA. **(e)** Correlation between replicates with DDA. **(f)** Correlation between replicates with DIA. **(g)** Ions measured in the orbitrap in MS2 scans with DDA & DIA. **(h)** Quantified difference in ions measured in the orbitrap for DDA & DIA. **(i)** Experimental workflow for evaluation of accuracy and precision of the method. **(j)** Boxplot of measured and theoretical ratios for yeast phosphopeptides with DDA & DIA. **(k)** Mean squared errors for DDA & DIA. **(l)** Receiver operating characteristic (ROC) curves for DDA & DIA.

However, even with the very fast and improved methodology, we seemed to have reached the limit for data-dependent acquisition (DDA) for phosphoproteomics with current instrumentation. Conversely, data-independent acquisition (DIA) can in principle overcome this limitation of sequential DDA by analyzing peptide ions in parallel. This is achieved by co-isolating co-eluting peptide ions in predefined mass windows, fragmenting them together and analyzing all the resulting fragment ions simultaneously. However, few attempts have been reported on applying DIA to large-scale (phospho)proteomics^11–15^. To address this problem we developed a PTM-specific workflow for peptide-centric DIA that combines the recovery rate of library-based extraction with high confidence site localization algorithm rivaling the current gold standard based on DDA.

Initially, we optimized the instrument settings for best DIA performance. Due to the co-fragmentation of multiple precursors with different charge-states in DIA, it is not possible to employ the charge-state dependent collision energy (CE) scaling of DDA. To identify the optimal collision energy settings for DIA, we recorded spectral libraries at four different normalized CE values and analyzed corresponding DIA runs with CE values fixed at charge-state of two. Analyzing the DIA files with the different spectral libraries revealed that the best compromise for maximizing identification of differently charged precursors was to record spectral libraries with NCE of 28 and analyze subsequent DIA runs with NCE of 25 at charge state 2 (**Supplementary Figure 1A**). Next, we optimized the overlap between adjacent mass windows in DIA for best quantification by quantifying precursors in overlapping regions in a DIA setup in which we systematically shifted the mass windows. From this analysis, we found that setting a fixed overlap of 1 Da between adjacent mass windows assured optimal quantification of precursors with m/z values at the edges of isolation windows (**Supplementary Figure 1B**). We also tested different DIA acquisition methods to find the optimal one for fast phosphoproteomics by changing scan cycle times using different mass window widths, number of windows and HCD resolution settings (**Supplementary Figure 1C**). All acquisition methods identified comparable numbers of phosphopeptides, which in DIA is defined as unique phosphorylated elution group precursors. However, the best quantitative performance judged by coefficient of variation (CV) between replica was achieved by the fastest scanning method employing 2 second cycle time with 48 mass windows of 14 Da widths using 15,000 resolution HCD fragmentation with maximum injection time of 22 ms (**Supplementary Figure 1D**).

Using this optimized DIA method with 15 min LC gradients, we identified almost three times as many elution group precursors and twice as many phosphopeptides compared to the number of DDA peptide-spectrum matches (PSMs) and phosphopeptides, respectively (**Figure 1B**, see methods section). In addition, the DIA raw files were also searched with directDIA (dDIA). In this approach, spectral libraries are generated directly by searching deconvoluted pseudo-MS/MS spectra from DIA data against a peptide database. For this process, Pulsar, the search engine in Spectronaut, applies the same search settings as DDA searches in MaxQuant. This library-independent dDIA strategy also worked well with twofold increase in precursors matched and 75% increase in phosphopeptides (**Figure 1B**). DIA further showed a significantly higher overlap of phosphopeptide identifications between replica compared to DDA (**Figure 1C and 1D**). Importantly, the quantitative reproducibility was better in DIA with correlation coefficient, R^2^ of 0.93 compared to R^2^ of 0.89 for DDA, even though DIA covered an additional order of magnitude of absolute precursor intensities (**Figure 1E and Figure 1F**). This is likely due to the fact that DIA makes more efficient use of the ion beam by sampling more ions in MS/MS mode compared to DDA throughout the entire LC-MS analysis (**Figure 1G**). Quantifying the difference reveals an approximate 6-fold higher fragment ion count measured in DIA compared to DDA mode (**Figure 1H**).

We next assessed the quantification accuracy and precision of the DIA and DDA approaches in a regulated phosphopeptide sub-population spiked into a complex sample background. For this purpose, we made use of a mixed species approach in which we diluted phosphopeptides enriched from yeast at different ratios into a fixed background of HeLa phosphopeptides and analyzed them by DDA, dDIA and DIA (**Figure 1I**). This strategy allows to assess how the acquisition methods quantify the expected ratios of the yeast phosphopeptides of 0.25:1, 0.5:1, 1. 5:1 and 2:1. As expected, dDIA and DIA were both able to quantify up to twice as many phosphopeptides as DDA (**Supplementary Figure 1E**). Based on box-plot analysis, all three methods very accurately estimated the expected ratios on median across all comparisons (**Figure 1J**). However, the interquartile ranges were significantly smaller for DIA compared to DDA and generally dropped at the higher loads (2:1) compared to the more dilute sample (0.25:1). This indicates that phosphopeptides of higher intensity are better quantified as expected. Conversely, the human phosphopeptides did not show regulation between the conditions (**Supplementary Figure 1F**).

To better assess quantification precision, we next calculated the mean squared error (MSE) as the sum of positive bias and variance for each method, which represent the quantification error in accuracy and precision, respectively (**Figure 1K**). Based on this, DIA yields the highest precision and highest accuracy at all ratios analyzed. DDA showed lower precision compared to dDIA at high intensity ratios (2:1), but better precision at low intensity ratios (0.25:1). To test if the accurate and precise quantification of DIA translated into a better identification of significant regulation, we used the significance d-score of the SAM test 16 to calculate true-positive-rates (TPR) and false-positive-rates (FPR) of the regulated phosphopeptides for the DDA and DIA approaches (**Figure 1L**). Plotting them against each other created a receiver operating characteristic (ROC) curve, in which best performing methods should achieve a TPR of 1 before increasing their FPR over 0. Focusing on the left part of the ROC curve plot, where the FPR is lowest, we see that DIA and dDIA shows the steepest TPR increase at all tested ratios compared to DDA.

### DIA-specific phosphorylation site localization algorithm

To compensate for the wider isolation windows applied in DIA, we developed a PTM localization algorithm for peptide centric analysis utilizing information not available in standard DDA data. This includes full isotopic patterns for fragment ions and the possibility of generating short elution chromatograms to correlate with the targeted precursor peak shape. The latter allows for systematic removal of any interfering fragment ions that one could not account for in DDA.

These two aspects are combined with additional scores based on fragment ion intensities and mass accuracy into a specific weighted score for each fragment, which is then used to calculate a specific site localization score (**Supplementary note 1**).

Briefly, during the standard DIA analysis, the algorithm starts out by detecting and classifying all potential peak groups for a peptide precursor in the library. For each candidate peak group modified peptides are enumerated into site candidates to represent all possible site combinations on-the-fly. Using the combined site information for a given peptide, the algorithm calculates and retains all unique fragment masses, which are then matched to the individual site candidates, whereby each individual fragment can be annotated as either confirming or refuting a specific site candidate (**Figure 2A**). A score is calculated for each site candidate using the individual fragment ion matches incorporating aspects of the feature, the mass accuracy and the XIC correlation. The final site scores are then calculated by summing all candidate scores that supported this site and subtracting all fragments that refute the site (**Figure 2B**). The developed DIA-specific PTM site localization score is finally calculated as a fractional probability score for each site combination compared in relation to all candidate scores. This approach is equivalent to the original PTM score site localization algorithm^2^ and the Andromeda score^17^ implemented in MaxQuant. The DIA-based PTM localization workflow does not require specially generated spectral libraries and is available in the Spectronaut software tool^7^ (v. 13.0.190309.20491).

**Figure 2.**
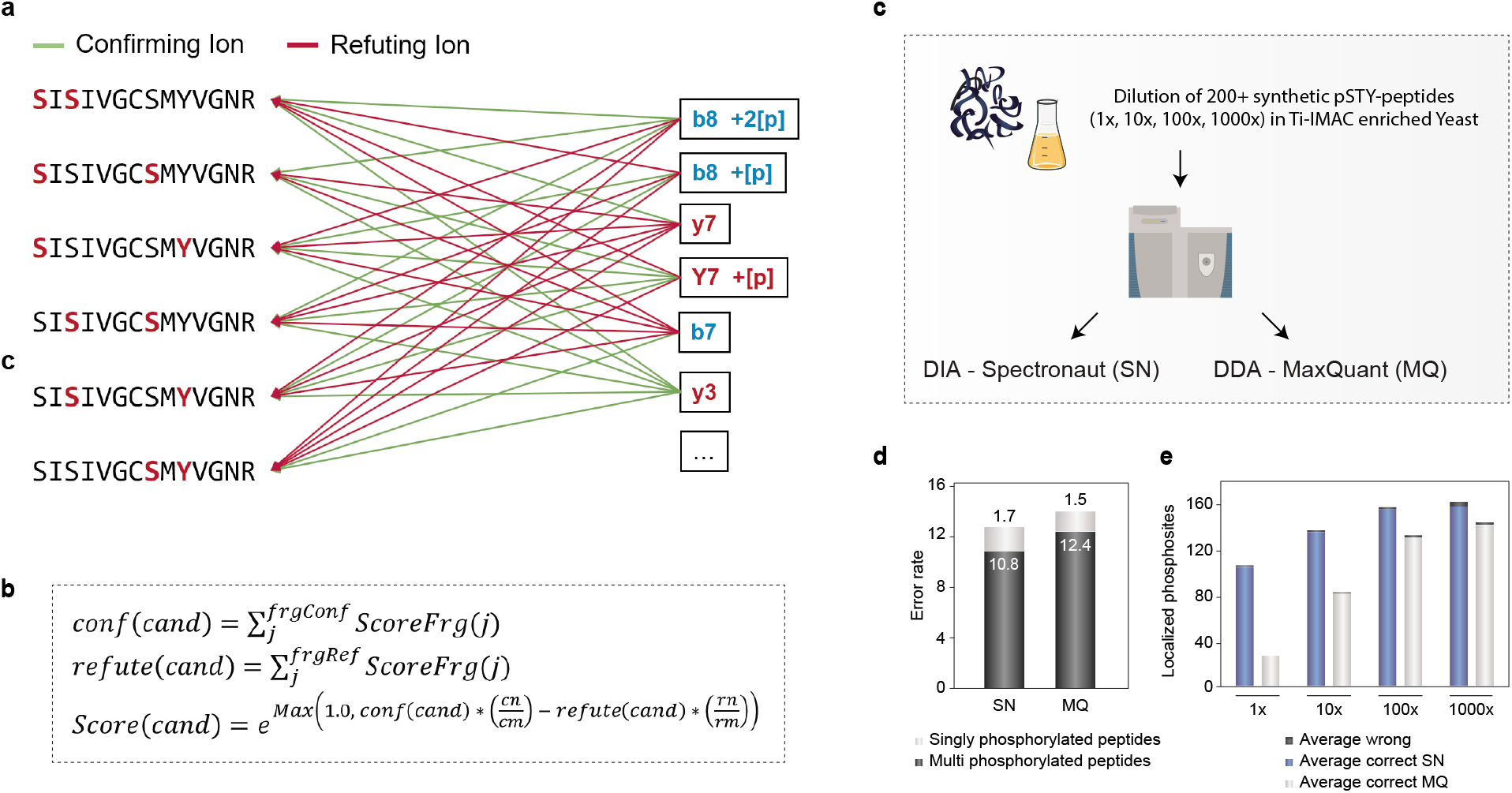
Challenges and solutions for phosphoDIA **(a)** Confirming and refuting fragments for site localization. **(b)** Calculation of final candidate score. **(c)** Workflow to evaluate localization algorithm with MaxQuant. **(d)** Error rates for phosphorylation site assignment in DDA and DIA. **(e)** Coverage of assigned phosphorylation site in a diluted yeast background for DDA & DIA.

To evaluate the performance of the PTM site localization algorithm and establish appropriate probability score cutoff values for accurate site localization, we initially tested it on a library of synthetic phosphopeptides with known site localization. This library consisted of two hundred human tryptic phosphopeptides frequently observed in large-scale phosphoproteomics experiments. They were spiked into a stable background of tryptic yeast phosphoproteome sample in different concentrations (1x, 10x, 100x, 1000x) and measured in triplicates using DDA and DIA (**Figure 2C**). The DDA raw files were analyzed with MaxQuant 18 using Phosphorylation (STY) as the only variable modification. The resulting phosphopeptide identifications were used to build a spectral library, which was then searched in peptide centric mode against the corresponding DIA raw files using Spectronaut with the PTM site localization algorithm implemented.

From the DDA files, we on average correctly localized 93.6 phosphorylation sites from singly phosphorylated peptides with 1.4% error rate on wrongly assigned sites (**Figure 2D**). We required at least 0.75 localization probability (Class I sites) as in previous analyses^2^, which in this case equates to an estimated FDR of 1.5%. Applying the same score cutoff of 0.75 for the DIA dataset results in correct identification and localization of 136.2 phosphorylation sites on average with 1.7% error rate of incorrectly assigned sites, indicating that site localization FDR at this cutoff value is comparable to that of DDA analyzed with MaxQuant but achieving higher site coverage in DIA. The site localization FDR when analyzing multiply phosphorylated peptides were somewhat higher but comparable between DDA and DIA with error rates of 12.4% and 10.8% respectively but the underlying statistics were limited due to fewer candidates. Notably, in opposition to DDA for which the number of synthetic phosphopeptide sites identified is significantly hampered at low dilutions, DIA maintained a relatively high identification rate across all dilutions, indicating that DIA outperformed DDA in sensitivity and dynamic range (**Figure 2E**). We also analyzed the DIA dataset of the synthetic phosphopeptides using dDIA without a spectral library and found that in this case we needed to apply a higher score probability cutoff of 0.99 to achieve error rates comparable to library-based DIA and DDA (**Supplementary Figure 2**). However even with this stringent cutoff, dDIA on average correctly identified and localized 128.8 phosphorylation sites, which was one-third more than DDA.

### Technical comparison of DDA and DIA in a biologically relevant setting

We next evaluated if the increased coverage of localized phosphopeptides in DIA over DDA translates into an advantage in a cell signaling study. For this purpose, we used EGF-stimulated retinal pigment epithelium (RPE1) cells treated with different MEK kinase inhibitors as a model system (**Figure 3A**). Briefly, cells were pretreated for 30 minutes with 0.5uM or 5uM of Cobimetinib, or 0.5uM or 5uM of PD-032591 prior to 10 minutes EGF stimulation, 10 minutes EGF only or untreated cells as a control. All conditions were prepared as biological triplicates, phosphopeptides from 200 ug of whole cell tryptic digests were enriched by Ti-IMAC and analyzed with 15 minute gradients by DDA and DIA.

**Figure 3.**
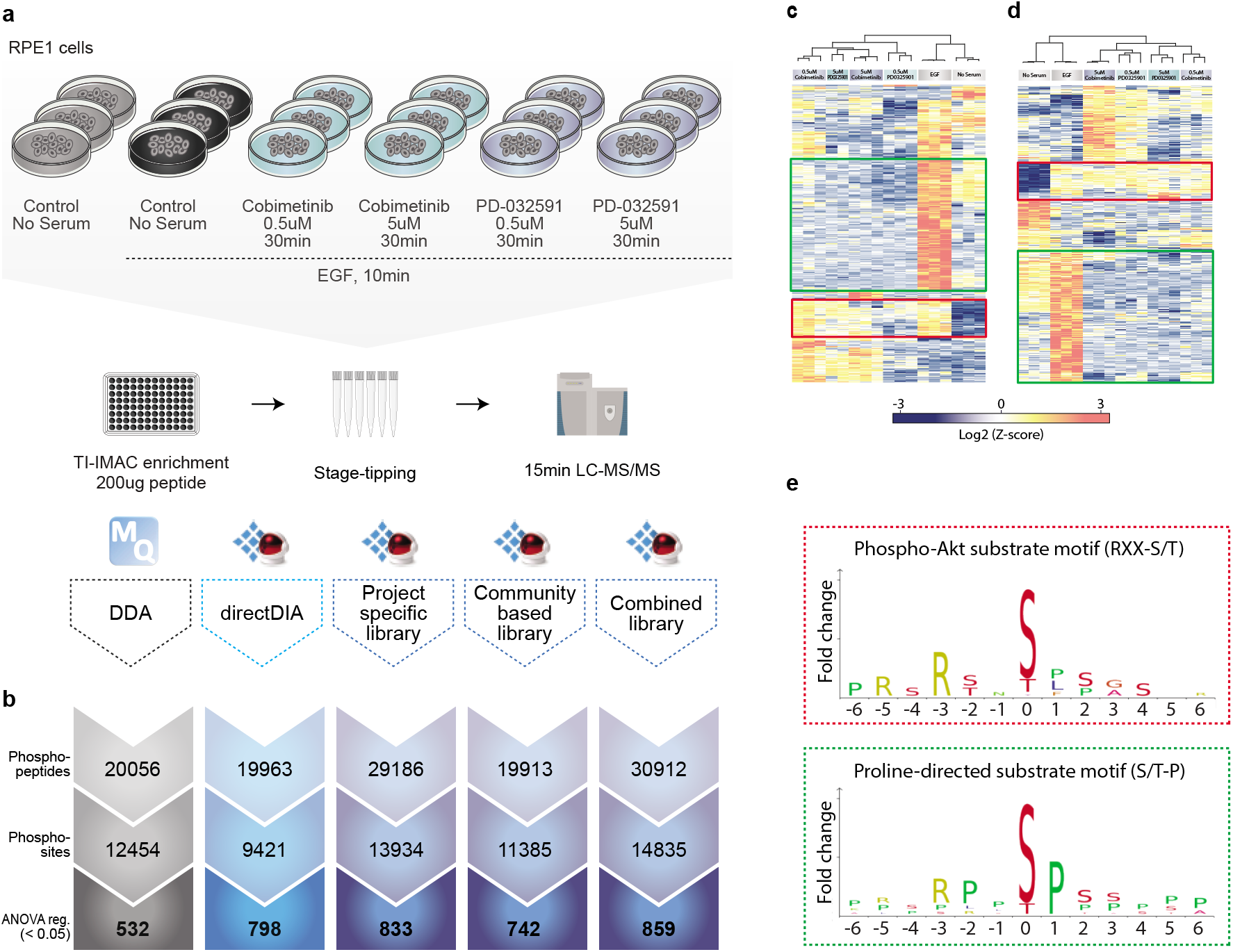
Technical comparison of DDA and different types of DIA in a biological setting **(a)** Experimental workflow **(b)** Overview of identified phosphopeptides, localized phosphosites and ANOVA regulated sites for the different methods **(c)** Heatmap of unsupervised clustering analysis of ANOVA regulated phosphosites for DDA workflow **(d)** and for DIA workflow with project specfic library **(e)** Linear sequence motif analysis for two selected clusters

For DIA, we tested the effect of using different spectral libraries and dDIA. We recorded a project-specific spectral library consisting of ~40,000 phosphorylation sites by deep phosphoproteome profiling^19^. We fractionated the same EGF-stimulated cells using massive offline high-pH reversed phase chromatography based fractionation combined with DDA analysis of individual fractions measured on the same 15 minute online LC-MS gradient as for the DIA analyses, which should be an ideal reference dataset (**Supplementary Figure 3A and Supplementary Figure 3B**). As an alternative to this, we created an even larger community-based spectral library of ~50,000 phosphorylation sites by combining two previous large-scale HeLa (phospho)proteome studies^20,21^ (**Supplementary Figure 3C**).

To facilitate efficient bioinformatics analysis of phosphoproteomics data, MaxQuant generates a site-level output table for each variable PTM that allows site-level statistical analysis. To compare DIA Spectronaut data to DDA MaxQuant data on a biological level, we developed a Perseus plugin that can convert a normal Spectronaut report into a site-level report (**Supplementary Note 1**). The plugin features a graphical interface and allows generation of MaxQuant-like site-level, PTM-localized peptide-level and “modification specific” peptide-level output.

With comparable data formats for both DDA and DIA, we first looked at the numbers of identified phosphopeptides and localized phosphorylation sites for each of the different methods, with more than ten thousand sites identified in all (**Figure 3B**). As expected, DDA with 20,056 phosphopeptides covering 12,454 sites identified less sites than DIA with project-specific spectral library with 29,186 phosphopeptides covering 13,934 sites. Conversely, the larger community-based library only identified 19,913 phosphopeptides and 11,385 sites, which is comparable to DDA. However, combining the two spectral libraries provides the best coverage with 30,912 phosphopeptides and 14,835 sites. Interestingly, dDIA yields similar number of phosphopeptides and sites as DDA and DIA with the community library.

To assess the biological effect of the different kinase inhibitor treatments on cellular phospho-signaling, we performed an analysis of variance (ANOVA) statistical test to identify significantly regulated sites. For this test, we only used phosphorylation site ratios quantified in all three biological replicates of at least one condition. Due to its lower precision and phosphopeptide coverage, DDA yielded 532 significantly regulated phosphorylation sites, whereas all DIA methods identified nearly twice the number of regulated sites (**Figure 3B**).

Next, we wanted to assess if and to what degree the regulated sites identified by DDA and DIA provided biological insights into EGF-dependent phospho-signaling in context of MEK inhibition as expected. To do this, we performed unsupervised hierarchical clustering of the DIA significant sites (**Figure 3C**) and the DDA significant sites (**Figure 3D**), which revealed an overall similar pattern of regulation between the different conditions. The pattern was the same for the other DIA methods (**Supplementary Figure 3D**). Linear sequence motif analysis of the EGF-upregulated phosphorylation sites that were largely unaffected by the kinase inhibitor treatment revealed that both DDA and DIA could correctly identify the EGF-dependent but MEK-independent AKT kinase substrate motif RxRxx[s/t]. Reassuringly, the ERK1/2 kinase substrate motif Px[s/t]P was significantly enriched among the MEK-dependent sites as expected (**Figure 3E**). This analysis demonstrated that DIA and DDA identified the same biology on localized phosphosite-level – AKT and ERK activation as the major signaling axes downstream of EGF – recapitulating known EGF receptor signaling as expected.

### Global analysis of fractional phosphorylation site stoichiometry from DIA data

Besides relative quantification of phosphorylation sites, it is valuable to determine their occupancy or absolute stoichiometry. A high fractional stoichiometry combined with dynamic regulation is a strong indication that the site is functional in the cellular context studied^1^. It is possible to determine the absolute occupancy or fractional stoichiometry of phosphorylation sites on a large scale by using ratios observed in both the phosphopeptide, its non-phosphorylated counterpart peptide and the respective protein between treatment conditions from SILAC data^22^ and TMT-multiplexed data^23,24^.

Based on our previous findings^24^, we reasoned that the high quantitative accuracy and the completeness of the DIA phosphoproteomics dataset should allow the extraction of stoichiometry from multiple conditions at the same time. We adapted a recently developed 3D multiple regression model (3DMM)-based approach based on TMT data to label-free DIA. The 3DMM approach integrates information of several experimental conditions and replicates into one stoichiometry model, which uses phosphopeptide-, non-phosphorylated peptide- and corresponding protein-intensities from any multiplexed quantification method. However even for DIA, data quantification of individual sites are not complete when analyzing many experimental conditions. To overcome this issue and retain as much quantitative information as possible, we combined peptide information based on the assumption of linear behavior between equally (non-)regulated peptides, which allowed us to extrapolate peptide intensities to fill in missing values. We implemented both the linear modeling approach and stoichiometry calculation into our Perseus plugin, which thus allows users without prior scripting experience to calculate PTM occupancies from LFQ data (**Supplementary Note 1**).

To benchmark the performance of the label-free 3DMM stoichiometry approach, we prepared a mixed species sample with fixed phosphopeptide stoichiometries (**Figure 4A**). A phosphopeptide-enriched tryptic yeast digest was split in two equal parts and one half was dephosphorylated using alkaline phosphatase. Mixing together phosphorylated and non-phosphorylated yeast peptides in fixed ratios into a HeLa phosphopeptide background yielded conditions of 1%, 10%, 50%, 90% and 99% phosphorylation site stoichiometry. Each sample was analyzed by DDA and DIA using 15 min LC-MS/MS. Boxplot analysis of the calculated stoichiometry values revealed that both DDA and DIA estimated the expected ratios closely on median across all comparisons. However, the interquartile ranges were significantly smaller for DIA compared to DDA especially at high stoichiometry values, indicating that higher intensity phosphopeptides are better quantified by DIA as expected (**Figure 4B**). To compare the quantification error in terms of accuracy and precision for the 3DMM data, we compared the mean squared errors (MSE) presenting the sum of positive bias and variance for the two methods, respectively, as described above. Based on this, DIA yields the highest precision and highest accuracy at all stoichiometry levels analyzed with the general trend that precision is lowest for highest values, whereas accuracy is more comparable across conditions (**Figure 4C**). Importantly, DIA was able to generate more than twice as many occupancy values (**Supplementary Figure 4A**), presumably due to its better quantification and higher phosphopeptide coverage (**Supplementary Figure 4B**).

**Figure 4.**
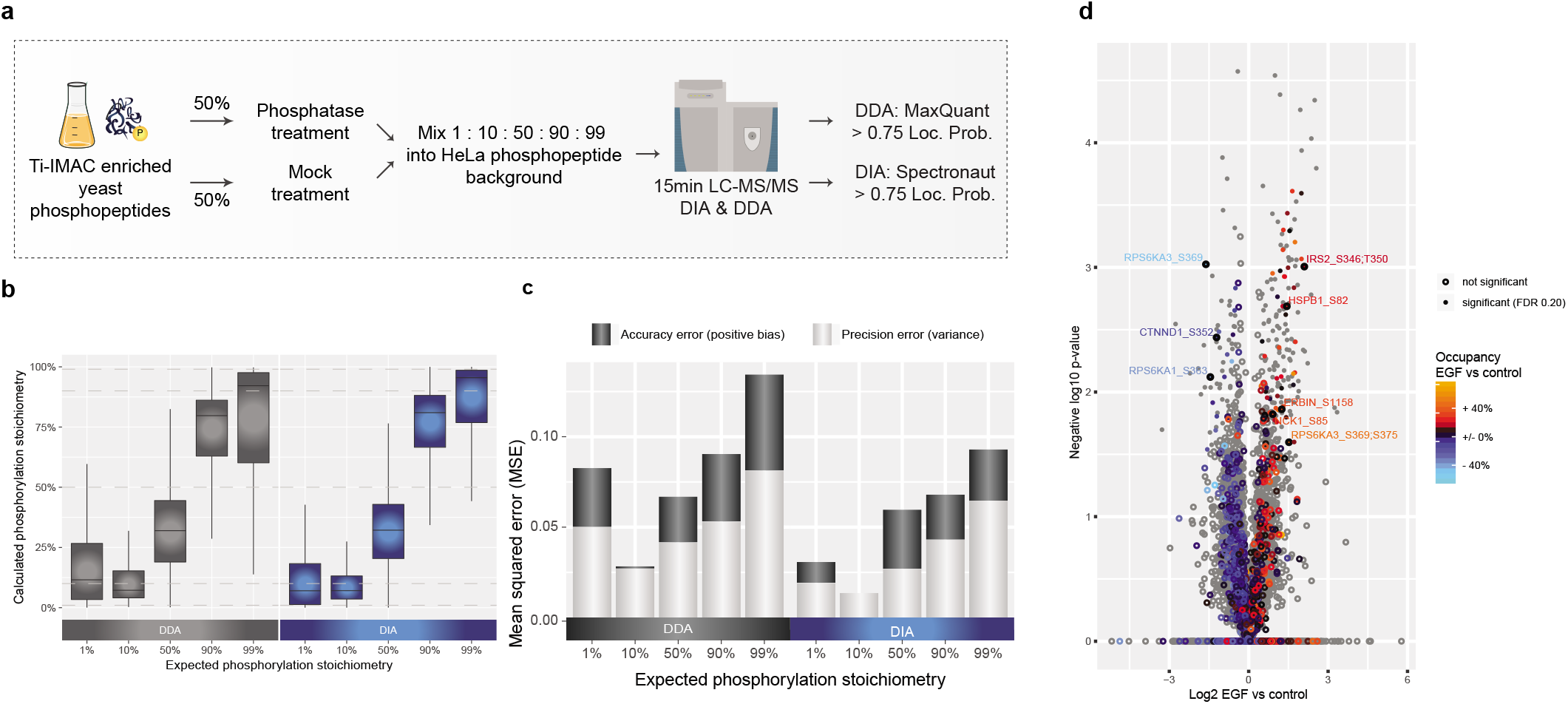
Stoichiometry benchmark **(a)** Experimental workflow for experiment with controlled ratios. **(b)** Boxplot of calculated phosphorylation stoichiometry with DDA & DIA. **(c)** Mean squared errors for calculated stoichiometries with DDA & DIA **(d)** Volcano plot analysis of calculated occupancies EGF vs. control

Our data indicates that DIA based stoichiometry estimation is possible with reasonable accuracy and we therefore applied it to the EGF-stimulated and kinase-inhibitor treated RPE1 cell phosphoproteome DIA dataset described above. Heatmap representation of the unsupervised hierarchical clustering of ANOVA significant phosphorylation site stoichiometries reflected the cellular conditions well with generally highest stoichiometry in EGF-stimulated samples and lowest in kinase inhibitor treated (**Supplementary Figure 4A**). This set of dynamic phosphorylation sites with high stoichiometry values was enriched in proteins associated with receptor tyrosine kinase (RTK) signaling according to the Reactome Pathway Database^25^. To pinpoint likely functional sites to prioritize for follow-up experiments, the global phosphorylation site stoichiometry measurements can be integrated with the corresponding site fold-changes and visualized as an extra layer of information in a Volcano plot exemplified by the comparison of EGF versus control samples (**Figure 4D**). Enrichment analysis among the significantly EGF-regulated site occupancies reveal strong overrepresentation of signaling by receptor tyrosine kinases and MAPK signaling pathways validating the known biology of the experiments (**Supplementary Figure 4B**).

### Large-scale DIA phosphoproteomics of hundreds of cell perturbations using a kinase inhibitor panel

To demonstrate the power and scalability of the rapid DIA-based site-specific phosphoproteomics workflow developed here, we applied it to identify phosphorylation site targets of the ten major protein kinases in the epidermal growth factor signaling pathway using a panel of thirty kinase inhibitors (**Figure 5A**). Briefly, EGF-stimulated RPE1 cells were pretreated with an inhibitor in two different concentrations (0.1uM and 1uM) in biological triplicates and each of the 186 samples was analyzed by 15 min LC-MS/MS using DIA. We quantified ~20,000 phosphopeptides across the 62 conditions in triplicates and performed ANOVA significance analysis on the log-transformed normalized intensities, which identified 1275 phosphorylation sites that were regulated in at least one condition. We visualized the regulated sites as a function of treatment by hierarchical clustering of the averaged phosphorylation site intensities per replica, which grouped likely substrates and targets according to the kinase inhibited (**Figure 5B**). From this analysis it is evident that EGF receptor inhibition (EGFRi) by all compounds worked well as it clusters with the untreated control samples. To verify that the inhibitors targeted the expected kinases, we performed a kinase motif enrichment analysis among each of the down-regulated phosphorylation site clusters for the individual kinase classes using a Fisher exact test. The overrepresented kinase motifs generally matched the expected kinase or their main established downstream kinase substrates (**Figure 5C**). For example, MEK inhibition leads to strong overrepresentation of ERK1/2 motif as expected, GSK3 inhibition down-regulates sites that conform to the known GSK3 motif, and mTOR inhibition down-regulated sites with the known mTOR substrate motif as well as substrate sites of its main downstream kinase p70S6K. To further validate the specificity of the kinase inhibitors we analyzed known substrates of the individual kinases and found good reproducibility in our dataset (**Figure 5D**).

**Figure 5.**
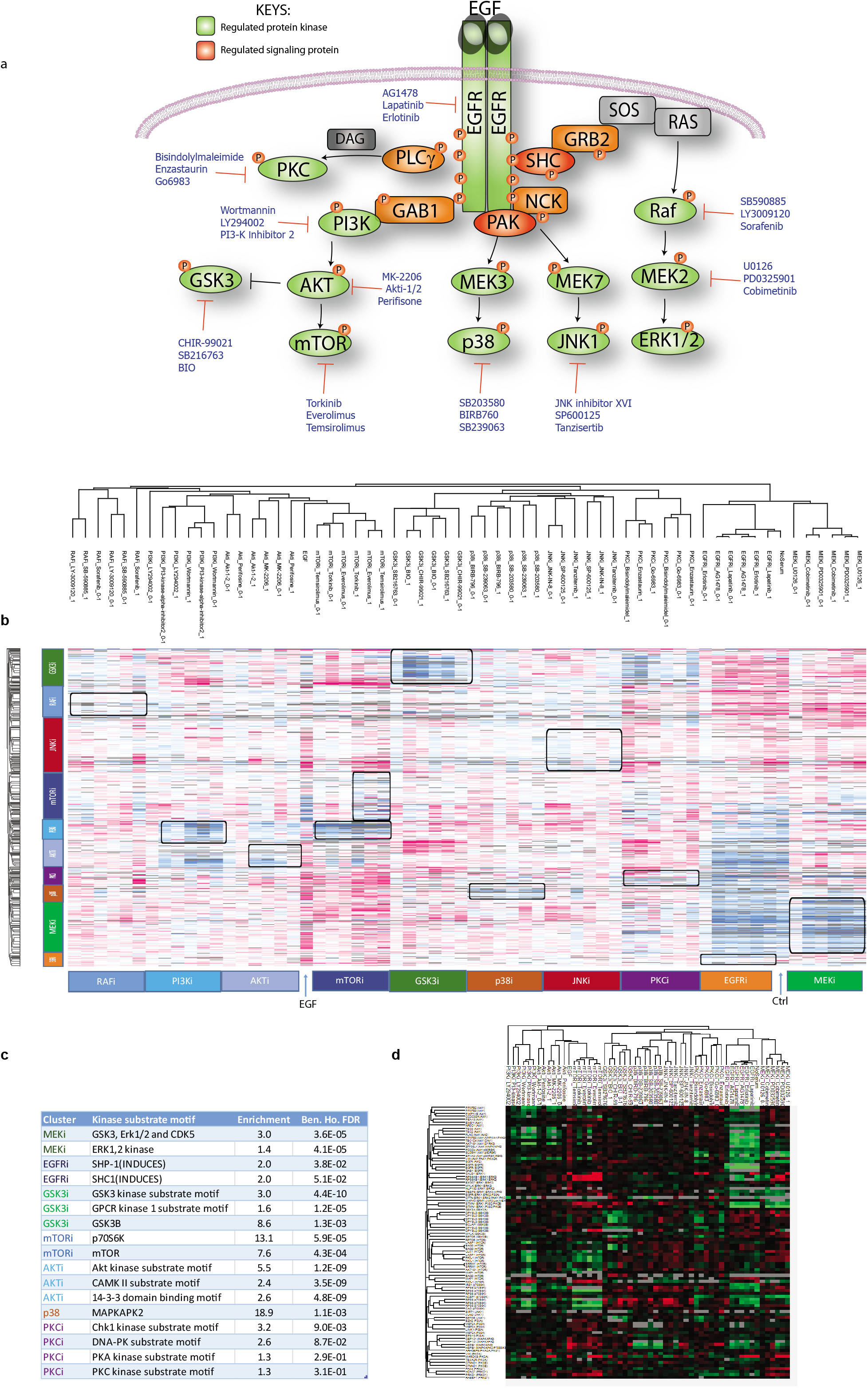
Kinase inhibitor screen **(a)** Experimental overview of kinase inhibitors used **(b)** Hierarchical clustering of averaged site intensities **(c)** Fisher Exact test of overrepresented kinase motifs **(d)** Clustering analysis of known substrated and individual kinases.

## DISCUSSION (Brief and focused)

Here we optimized a streamlined phosphoproteomics workflow based on data-independent acquisition (DIA) and developed a PTM localization site algorithm as part of the DIA computational pipeline, which we benchmarked against state-of-the-art DDA-based phosphoproteomics. By analyzing a library of synthetic phosphopeptides, controlled mixed-species phosphoproteomes with technical replication and EGF-stimulated cells in combination with kinase inhibitors with biological replication, we show the robustness, specificity, and high quality of the DIA-based phosphoproteomes. Quantitatively we demonstrate that we can achieve significant greater depth than any previous DIA-based phosphoproteome reported. Moreover, to our knowledge, we present the first systematic analysis of kinase inhibitor characterization by phosphoproteomics using label-free quantification.

More generally, the methodological approach we have developed represents a strategy to quantitatively profile hundreds of phopshoproteomes in a few days from low amount of starting material. While it has been shown previously that sensitive phosphoproteomics starting from 200 ug or less^26^ is feasible, this study represents an important advancement in that we use sensitive phosphoproteomics to analyze cellular signaling networks much faster and with greater depth. Furthermore, we demonstrate that stoichiometry calculation using LFQ is principally feasible, which has the potential to help in identifying functionally relevant phosphorylation sites in the future.

There are still limitations to the optimal DIA-based phosphoproteomics workflow. We achieve the best coverage and quantification when using tailor-made project specific spectral library for the DIA analyses, but this requires some effort and may not always be possible. However, predicting peptide retention time and MS/MS spectra^27,28^ might circumvent the necessity of recording spectral libraries for DIA in future. Unfortunately, the current prediction tools are not yet developed for phosphoproteomics.

Alternatively, library-free approaches such as dDIA look very promising for the future. Even though it takes a significant hit in the number of identified phosphopeptides, it is much easier to implement. Importantly, dDIA also may overcome issues with standard DIA, where rare or low abundant phosphorylation sites may get diluted out during library generation. Moreover, we believe that there is still room for improvements in the mass spectrometric technology. Although we show that DIA analyzes about 6-fold more ions in MS/MS mode than DDA, we estimate that by using 48 DIA-windows we are still maximally sampling a few percent of the ion beam at best.

Mass spectrometers with higher duty cycles such as the timsTOF pro^29^ may be an even better fit for DIA-based phosphoproteomics in the future. Looking forward, the methodological and computational framework outlined here may be applicable to a clinical setting for example in cancer, where sensitive phosphoproteomics profiling of individual patient tumors may aid precision medicine in the future.

## ONLINE METHODS

(all technical details necessary for the independent reproduction of the methodology, without referring to a chain of bibliographical references)

### Human cell culture and lysis

Human epithelial cervix carcinoma HeLa cells (female) and human retinal pigment epithelial RPE1 cells (female) immortalized with hTERT were purchased from ATCC and cultured in DMEM (Gibco, Invitrogen) supplemented with 10% fetal bovine serum (Gibco, Invitrogen), 100U/mL penicillin (Gibco, Invitrogen), 100ug/mL streptomycin (Gibco, Invitrogen) at 37 °C in a humidified incubator with 5% CO2. Cells were harvested at approximately 80% confluency by washing twice with PBS and (Gibco, Invitrogen) and subsequently adding boiling GdmCl lysis buffer (6M guanidine hydrochloride, 5 mM tris(2-carboxyethyl)phosphine, 10 mM chloroacetamide, 100 mM Tris pH 8.5 directly to the plate. Cells were collected by scraping the plate and boiled for 10min at 95°C followed by micro tip sonication.

### Yeast cell culture and lysis

BY4742 wild-type cells were grown in YEPD medium (2% bacto peptone, 1% yeast extract, 2% dextrose/glucose; sterile-filtered before use) at 30°C and 200 rpm rotation in overnight culture. Day culture was inoculated at OD_600 of 0.1 and harvest hours later when the OD_600 surpassed ~0.8. Yeast cells were spun down (4000g 5min) and washed with ice cold PBS. The pellet was resuspended in yeast lysis buffer (20ml per 1l OD_600 1; 75mM Tris pH8, 75mM NaCl, 1mM EDTA, 1 complete mini protease inhibitor cocktail tablet per 10ml, 5 mM sodium fluoride, 1 mM sodium orthovanadate, 5 mM β-glycerol phosphate) and dropped in droplets out of pipette into liquid nitrogen. Frozen droplets were ground in a MM400 ball mill (Retsch) for 3 min at 25 Hz. Frozen yeast powder was then mixed with 1% Triton X-100 and 0.5% SDS and incubated rolling at 4°C until thawed. Yeast lysate was spun down (16,000rpm 4°C 5min). The supernatant was transferred into –80°C acetone to a final acetone concentration of 80% v/v and incubated at −20°C for 2h. Precipitated proteins were spun down and resuspended in GdmCl lysis buffer (6M guanidine hydrochloride, 5 mM tris(2-carboxyethyl)phosphine, 10 mM chloroacetamide, 100 mM Tris pH 8.5, 1 complete mini protease inhibitor cocktail tablet per 10ml, 5 mM sodium fluoride, 1 mM sodium orthovanadate, 5 mM β-glycerol phosphate). Protein pellets were resuspended by sonication (Sonics & Materials, VCX 130; 1s on, 1s off, 80% amplitude) and boiled for 10min at 95°C.

### Protein digestion

Protein concentration was estimated by BCA assay (Pierce) and the lysates were digested with Lys-C (Wako) in an enzyme/protein ratio of 1:100 (w/w) for 1h followed by a threefold dilution with 25mM Tris, pH 8.5 to 2M GdmCl and further digested overnight with trypsin (Sigma Aldrich) 1:100 (w:w). Protease activity was quenched by acidification with trifluoroacetic acid (TFA) to a final concentration of approximately 1% and the resulting peptide mixture was concentrated using reversed-phase Sep-Pak C18 cartridge (Waters). Peptides were eluted off the Sep-Pak with 2mL 40% acetonitrile(ACN) followed by 2mL 60% ACN. The ACN was removed by vacuum centrifugation at 60°C and the final concentration was estimated by measuring absorbance at 280 nm on a NanoDrop 2000C, Thermo Scientific).

### Phosphopeptide enrichment

200ug of peptides were enriched for phosphopeptides using Ti-IMAC magnetic beads (Resyn Biosciences). Enrichments were carried out in protein LoBind 96-well plates (Eppendorf). The plates were mixed using 1300 rpm (Heidolph Titramax 1000, #544-12200-00) and separated in the 96-well plate using a magnetic stand (Thermo Scientific, # AM10027). Ti-IMAC beads were equilibrated twice in 200 μl 70% Ethanol followed by once in 100 μl 1% Ammonia and three times in loading buffer (80% acetonitrile, 1M glycolic acid, 5% TFA). 200 μg peptide mixture was mixed with equal amount of loading buffer and 500 μg Ti-IMAC beads (25 μl) were added and the solution was mixed for 20min. The beads were separated on a magnetic stand for 20s and the supernatant was removed using gel-loader tip connected to vacuum. 200 μl of loading buffer was added and beads were mixed for 2min followed by two washes with wash buffer 1 (80% acetonitrile, 1% TFA) and once with wash buffer 2 (10% acetonitrile, 0.2% TFA). Phosphopeptides were eluted in three rounds with 80 μl 1% Ammonia for 20 min. Transfer the supernatant to a clean plate and acidify with TFA. Speedvac solution and desalt using C18 StageTips and store at 4°C until MS analysis.

### Preparation of spectral libraries

RPE1 cells were stimulated with 125 ng/mL EGF (Chromotek) for 3 and 10 min followed by lysis and digestion as described before. Ultimate 3000 ultra-high performance liguid chromatography (UHPLC) (Dionex, Sunnyvale, CA, USA), in conjunction with high-pH reversed-phase chromatography, was used to separate and fractionate tryptic peptides. Peptides were separated using a high-pH-compatible 250 3 4.6 mm C18 Waters BEH X-Bridge peptide separation technology (PST) 3.6 mM or Phenomenex Cell Reports 22, 2784–2796, March 6, 2018 2793 Kinetex Evo 2.6 mM (Torrance, CA USA) column with identical dimensions. Basic conditions were achieved by running buffer C (50 mM ammonium hydroxide) constantly at 10% (100 mL/min, 2.5 mM ammonium bicarbonate final). A 60 min fractionation and collection gradient was achieved using buffer A (Milli-Q H2O) and buffer B (acetonitrile), and fractions were collected. From 3mg, 10 fractions were collected, while 10 mg was fractionated and collected in 46 fractions without any concatenation. Running at a constant 1 mL/min, the gradient was increased from 5% to 25% buffer B in 50 min and further increased to 70% buffer B in 5 min, where it was held for another 5 min. At this point, the fraction collection was stopped. Each fraction was dried in a speedvac and reconstituted in phopsho loading buffer and phosphopeptides were enriched as described above. Each fraction was analyzed individually with LC-MS/MS settings as described below.

### Stoichiometry

Phosphopeptides enriched from yeast and HeLa cells as described above were each split 50:50. One half was dephosphorylated using rAPid alkaline phosphatase (Sigma-Aldrich; 1 ul per 2mg protein starting material) at 37°C 750rpm shaking overnight, while the other half was mock treated using water. Both samples were incubated for 10min at 85°C to inactivate the phosphatase, and then purified on C18 StageTips^30^. Yeast phosphopeptides were mixed at 0. 1%, 1%, 10%, 50%, 90%, 99% and 99.9% into dephosphorylated peptides at 99.9%, 99%, 90%, 50%, 10%, 1% and 0.1%, with the total amount of peptides per condition corresponding to Ti-IMAC enrichment from 200ug yeast starting material. Mixtures were then added into a 1:1 HeLa mixture of phosphopeptides and dephosphorylated peptides, each corresponding to a total amount of peptides per condition corresponding to Ti-IMAC enrichment from 100ug HeLa starting material. Samples were then measured with the DDA and optimized DIA method as described below.

### Synthetic phosphopeptide experiment

Synthetic phosphopeptides were purchased from JPT (SpikeMix PTM-kit 52 1001098; SpikeMix PTM-kit 54 1001100) and Sigma-Aldrich (MS PhosphoMix 1 Light MSP1L, MS PhosphoMix 2 Light MSP2L, MS PhosphoMix 3 Light MSP3L) and resuspended according to the manufacturer’s instructions. Synthetic phosphopeptides and yeast phosphopeptides as described above were mixed in different ratios to mimic complex phosphopeptide mixtures (**Supplementary Table 1**). Samples were measured in DIA and DDA mode in technical triplicates each. For DIA measurement, the optimized method described below with the LC-gradient scaled to 35min was used. For DDA measurement, the same LC gradient was measured with MS1 resolution 60,000, MS1 AGC target 3e6, MS1 max IT 45ms, scan range 350-1400 m/z, MS2 resolution 30,000, MS2 AGC target 1e5, MS2 max IT 54ms, MS2 top6, MS2 isolation window 1.3 m/z, MS2 scan range 200-2000 m/z, MS2 NCE 28% and MS2 dynamic exclusion 30sec. DIA data was searched using a spectral library generated from the DDA files, searched with MaxQuant and generated by Spectronaut as described below.

### Nanoflow LC-MS/MS

The peptides concentrated in a speedvac and volume were adjusted to 7 μl in loading buffer (5% ACN and 0.1% TFA) prior to autosampling. An in-house packed 15 cm, 75 μm ID capillary column with 1.9 μm Reprosil-Pur C18 beads (Dr. Maisch, Ammerbuch, Germany) was used. An EASY-nLC 1200 system (Thermo Fisher Scientific, San Jose, CA) was and the column temperature was maintained at 40 °C using an integrated column oven (PRSO-V1, Sonation, Biberach, Germany) interfaced online with the mass spectrometer. Formic acid (FA) 0.1% was used to buffer the pH in the two running buffers used. The total gradient time was 19 min and went from 8 to 24% acetonitrile (ACN) in 12.5 min, followed by 2.5 min to 36%. This was followed by a washout by a 1/2 min increase to 64% ACN, which was kept for 3.5 min. Flow rate was kept at 350 nL/min. Re-equilibration was done in parallel with sample pickup and prior to loading with a minimum requirement of 1 μL of 0.1% FA buffer at a pressure of 800 bar.

Spray voltage was set to 2 kV, funnel RF level at 40, and heated capillary at 275 °C. For DDA experiments full MS resolutions were set to 60,000 at m/z 200 and full MS AGC target was 3E6 with an IT of 25 ms. Mass range was set to 350-1400. AGC target value for fragment spectra was set at 1E5 with a resolution of 15,000 and injection times of 22ms and Top12. Intensity threshold was kept at 2E5. Isolation width was set at 1.3 m/z and a fixed first mass of 100 m/z was used. Normalized collision energy was set at 28%. For DIA experiments full MS resolutions were set to 120,000 at m/z 200 and full MS AGC target was 3E6 with an IT of 45 ms. Mass range was set to 350-1400. AGC target value for fragment spectra was set at 3E6. 48 windows of 14 Da were used with an overlap of 1 Da. Resolution was set to 15,000 and IT to 22 ms.Normalized collision energy was set at 25%. All data were acquired in profile mode using positive polarity and peptide match was set to off, and isotope exclusion was on.

### Raw data processing

DDA files were processed using MaxQuant (1.6.5.0) with default settings. Carbamidomethyl (C) was set as fixed modifications. Oxidation (M), Acetyl (Protein N-term), Phospho (STY), Deamidation (NQ) and Gln->pyro-Glu were set as variable modifications. Reference FASTA files for human and *S. cerevisiae* were downloaded from Uniprot on 15th of March 2018. Spectral libraries were built from MaxQuant DDA search results using Spectronaut Professional x64 (13.0.190309.20491)^7^ with default settings, but “Best N Fragments per Peptide Max” set to 25 instead of 6. The yeast phosphopeptide library was generated from yeast phosphopeptides fractionated into 12 and 46 fractions. We used an Ultimate 3000 HPLC system (Dionex) with a Waters Acquity CSH C18 1.7 μm 1×150 mm column on operating at a flow rate of 30 μl/min with two buffer lines as previously described^21^.

DIA files were processed using Spectronaut with default settings, with “PTM localization” activated and probability cutoff set to 0 (for peptide-level analysis of dilution benchmark) or 0.75, “Data filtering” set to “Qvalue” and “Normalization Strategy” set to “Local Normalization”. Unless otherwise stated, experiments containing yeast peptides were searched using the yeast phosphopeptide library, those containing human peptides using the human phosphopeptide library, and those containing both with both. For the DIA stoichiometry benchmark (**Figure 4A-C**), DIA files were additionally searched with a library generated from the DDA runs of the same experiment in order to have non-phospho peptides represented in the library as well. The synthetic phosphopeptide benchmark (**Figure 2**) was searched with a library generated from the DDA runs of the same experiment.

Direct DIA search was performed in Spectronaut using default settings with the same PTMs as defined in MaxQuant DDA searches, “Data filtering” set to “Qvalue” and “Normalization Strategy” set to “Local Normalization”.

Transformation of the Spectronaut normal report and calculation of stoichiometry values was performed using a custom coded plugin “Peptide Collapse” in Perseus (1.6.5.0). The plugin was created using Microsoft Visual Studio 2017 (15.6.3) and requires Perseus and R (minimum version 3.6.0) to run. Detailed information on how to install and use the plugin and the calculations it performs are listed in **Supplementary Note 1**.

### Bioinformatics analysis

Most of the data analysis was performed using custom scripts in R (64bit version 3.6.0) with packages data.table (1.12.2), bit64 (0.9-7), doParallel (1.0.14), stringr (1.4.0), ggplot2 (3.1.1), qplot (3.0.1.1), limma (3.40.0), samr (3.0), magrittr (1.5), scales (1.0.0), XML (3.98-1.19), PerseusR (0.3.4), Biostrings (2.52.0) and MASS (7.3-51.4). ANOVA testing was performed in Perseus (1.6.5.0) and KEGG/reactome term enrichment was performed using the STRING app (1.4.2) in Cytoscape (3.7.0).

For the dilution benchmark experiment (**Figure 1I-L**), Spectronaut reports were transformed into “modification specific peptide”-like reports using the plugin peptide collapse in Perseus, with EG.PTMAssayProbability as grouping column, localization cutoff 0 and same variable PTMs as listed above. For the stoichiometry benchmark experiment (**Figure 4A-C**) and the kinase inhibitor stoichiometry experiment (**Figure 4D**), stoichiometry values on “target PTM peptide-level” were calculated for both DDA and DIA data using the plugin peptide collapse in Perseus, with grouping columns “Modified sequence” and “EG.PTMLocalizationProbabilities”, respectively. Localization cutoff was set to 0.75 for both and same variable PTMs as listed above.

For the LFQ dilution and stoichiometry benchmark experiment, Spectronaut intensities were “de-normalized” by dividing reported intensity values by their normalization factors. For the stoichiometry benchmark experiment, DDA and DIA intensities were then quantile-normalized. Boxplots were created with boxes marking the first and third quartile, a dash the median, and whiskers the minimum/maximum value within 1.5 interquartile range. Outliers are not displayed. SAM testing was performed using default settings (s0 =0.1, FDR = 0.20). ANOVA testing of stoichiometry values was performed using s0 = 0.01 and FDR = 0.20. Enrichment of KEGG/reactome terms was performed in the Cytoscape STRING app using default settings.

### Code availability

Custom R code for the data analysis and the Perseus plugin “Peptide Collapse” are available upon request.

## ACKNOWLEDGEMENTS

We would like to acknowledge Jan Rudolph for his help in coding and designing the Perseus peptide collapse plugin. Work at The Novo Nordisk Foundation Center for Protein Research (CPR) is funded in part by a generous donation from the Novo Nordisk Foundation (Grant number NNF14CC0001). The proteomics technology developments applied was part of a project that has received funding from the European Union’s Horizon 2020 research and innovation programme under grant agreements: MSmed-686547, EPIC-XS-823839 and ERC synergy grant 810057-HighResCells.

---

**Supplementary Figure 1.**
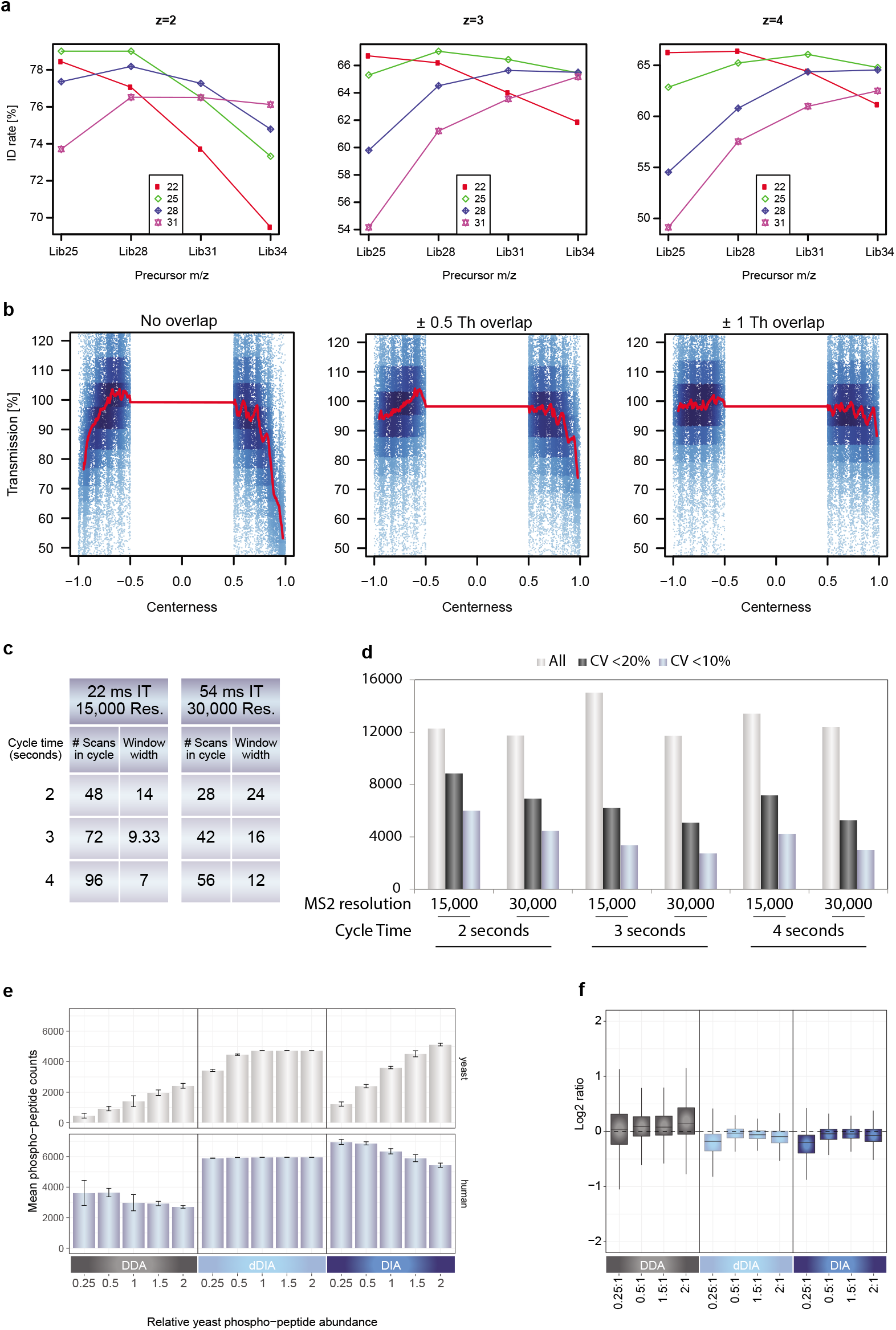
High-throughput and sensitive phosphoproteomics for DDA & DIA – identification and quantification **(a)** Comparison of optimal NCE for spectral library generation and DIA runs **(b)** Comparison of overlap between mass windows **(c)** Schematic overview of methods used for cycle time optimization **(d)** Average identifications of phosphopeptides and the number of phosphopeptides with CVs below defined thresholds **(e)** Number of HeLa and yeast phosphopeptides measured with DDA, DIA and dDIA **(f)** Boxplot of measured and theoretical ratios for hela phosphopeptides with DDA & DIA.

**Supplementary Figure 2.**
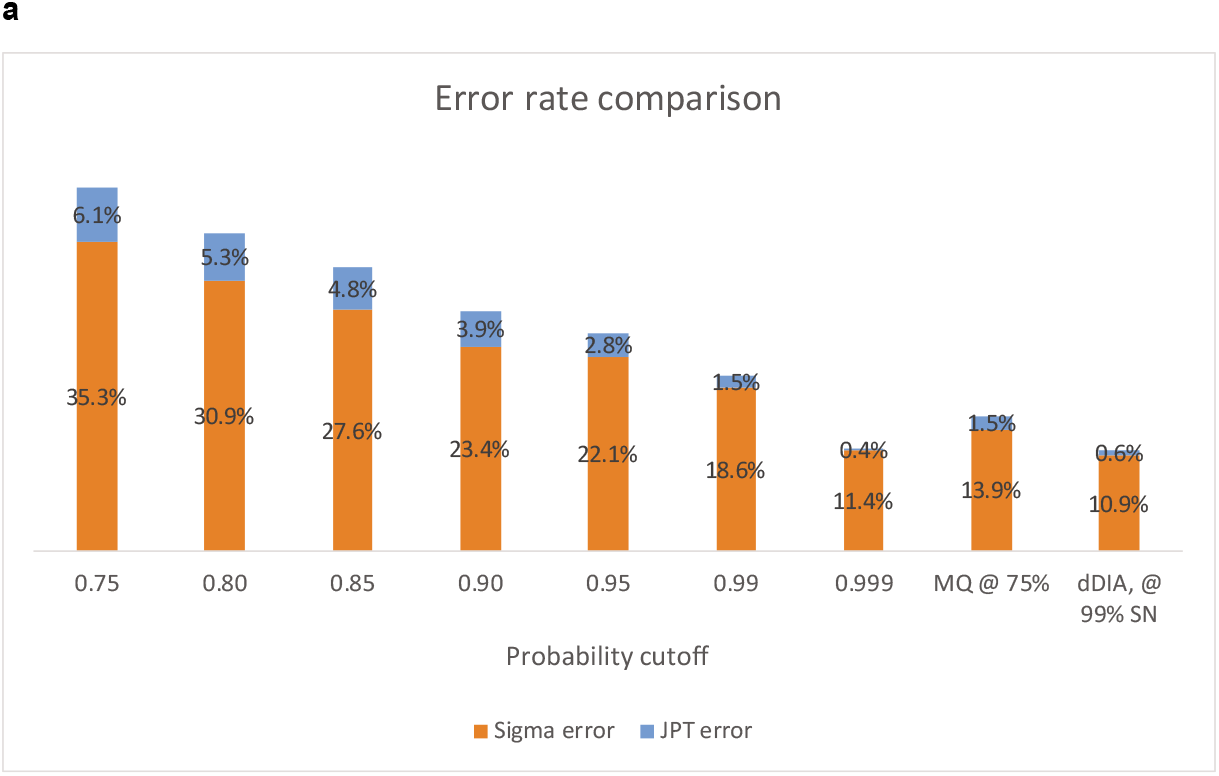

**Supplementary Figure 3.**
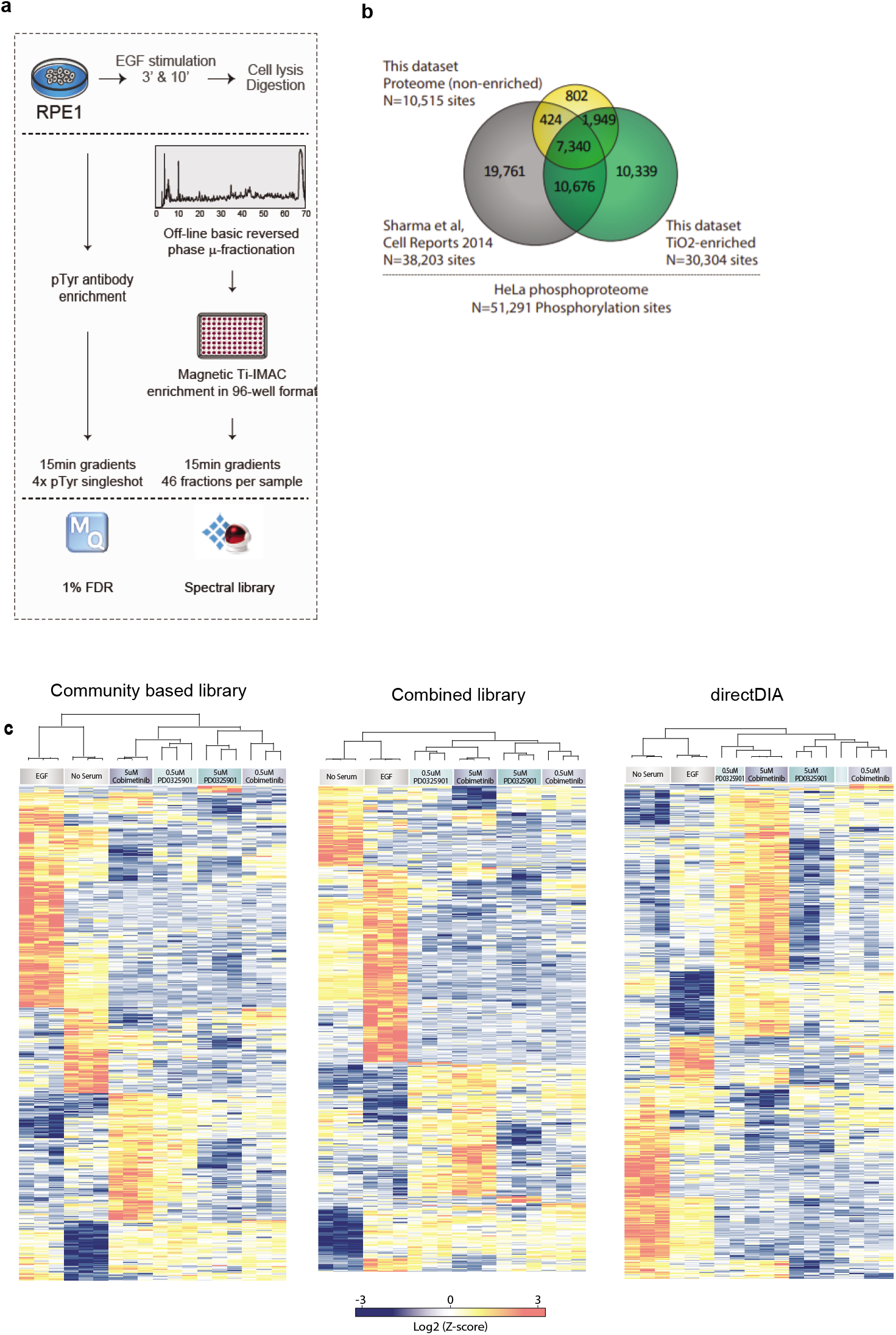
Technical comparison of DDA and different types of DIA in a biological setting **(a)** Experimental workflow for building spectral library **(b)** Overview of community based library **c)** Heatmap of unsupervised clustering analysis of ANOVA regulated phosphosites for DIA workflows with community based library, combined library and directDIA.

**Supplementary Figure 4.**
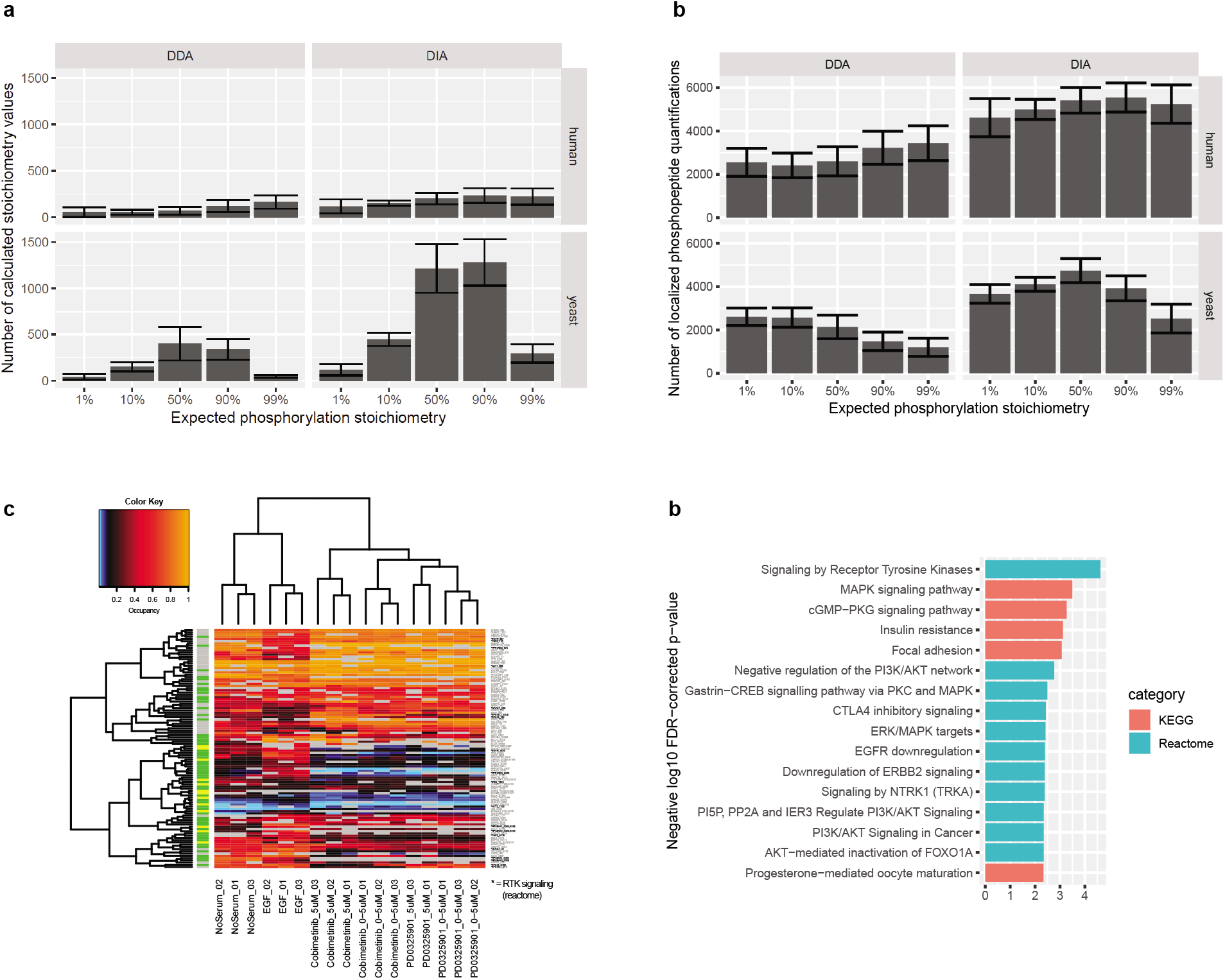
Stoichiometry benchmark **(a)** Number of clculated stoichiometry values **(b)** Number of measured peptide quantifications **(c)** Heatmap of ANOVA, FDR 0.20 regulated occupancies **(d)** Enrichment analysis of ANOVA significant occupancies

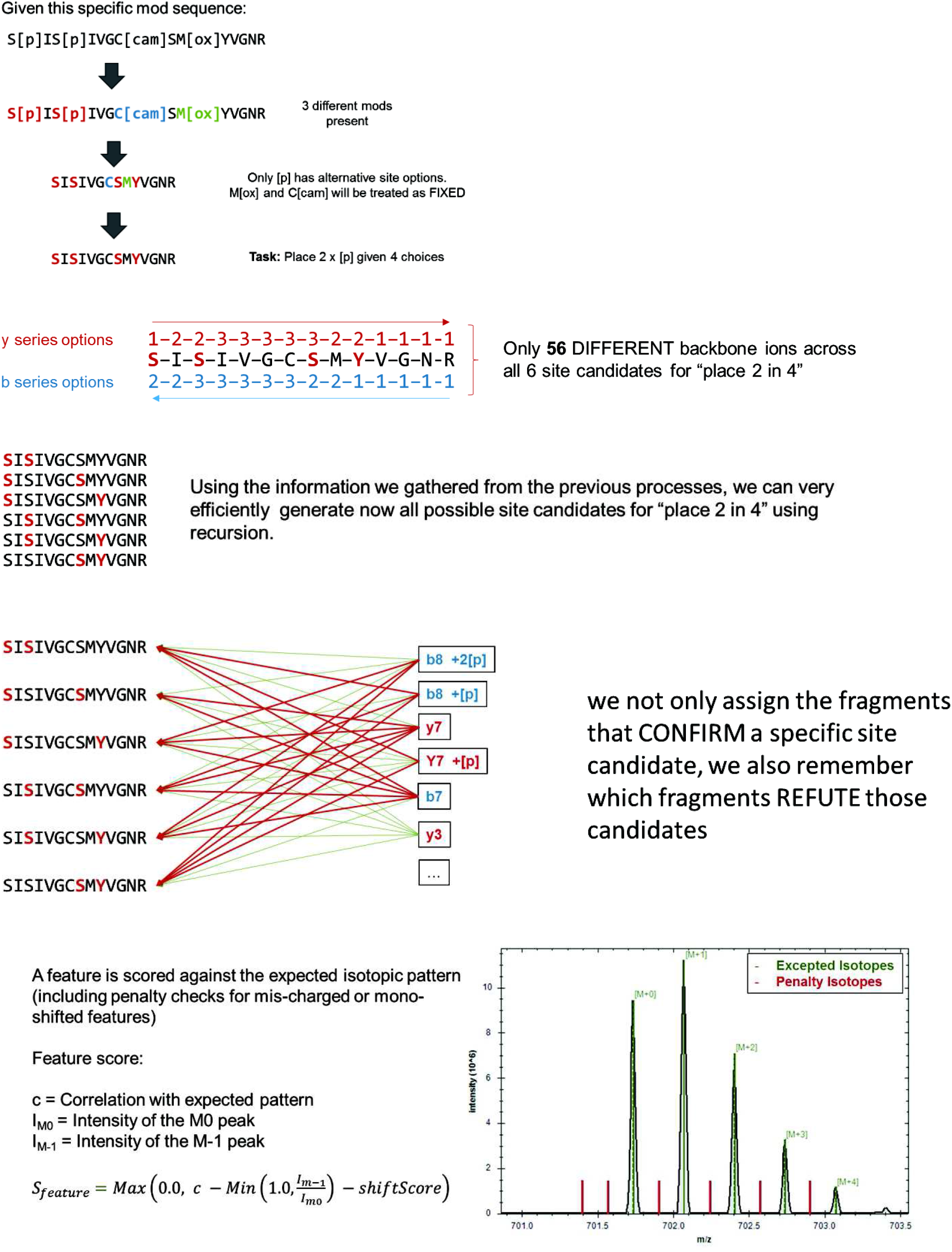

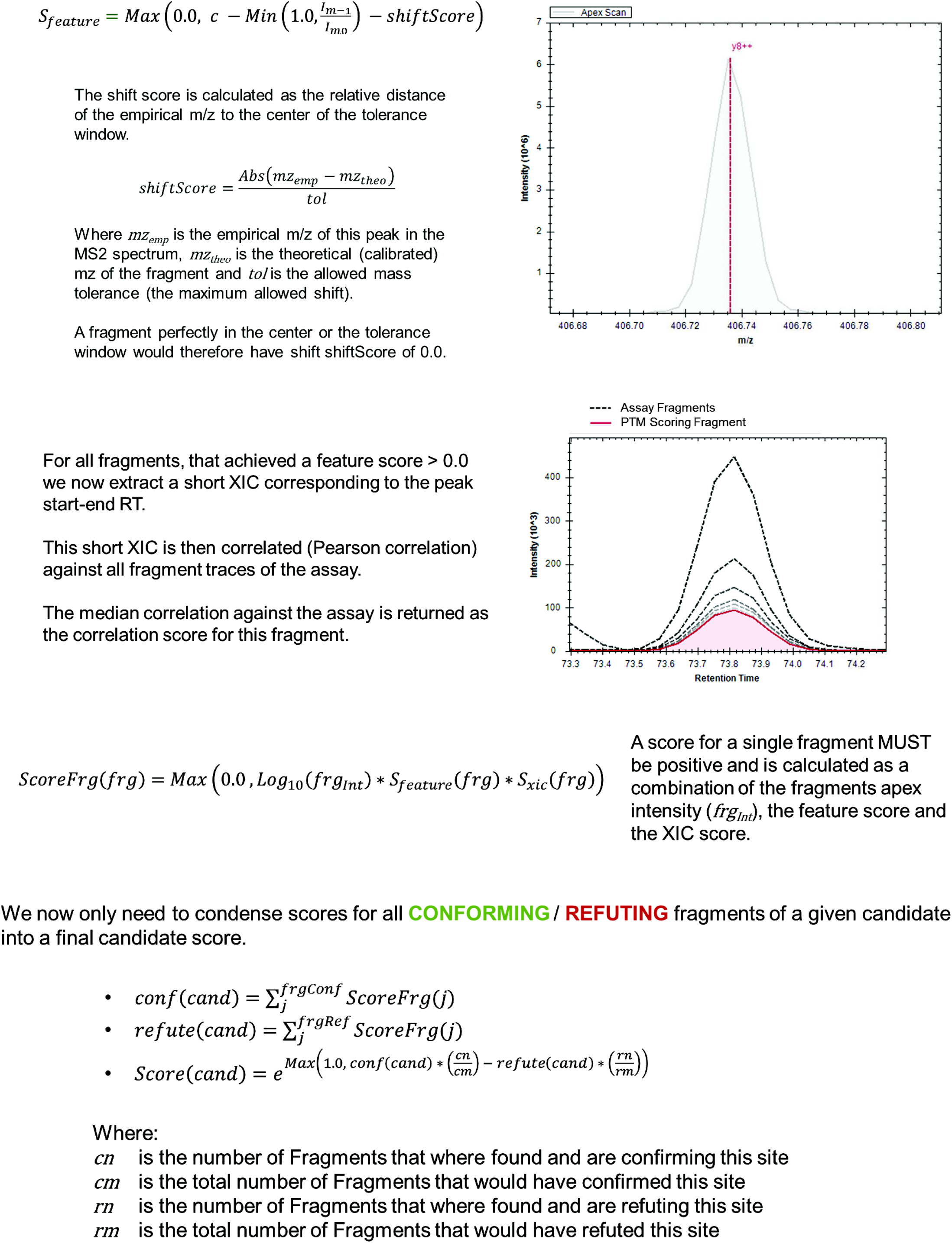
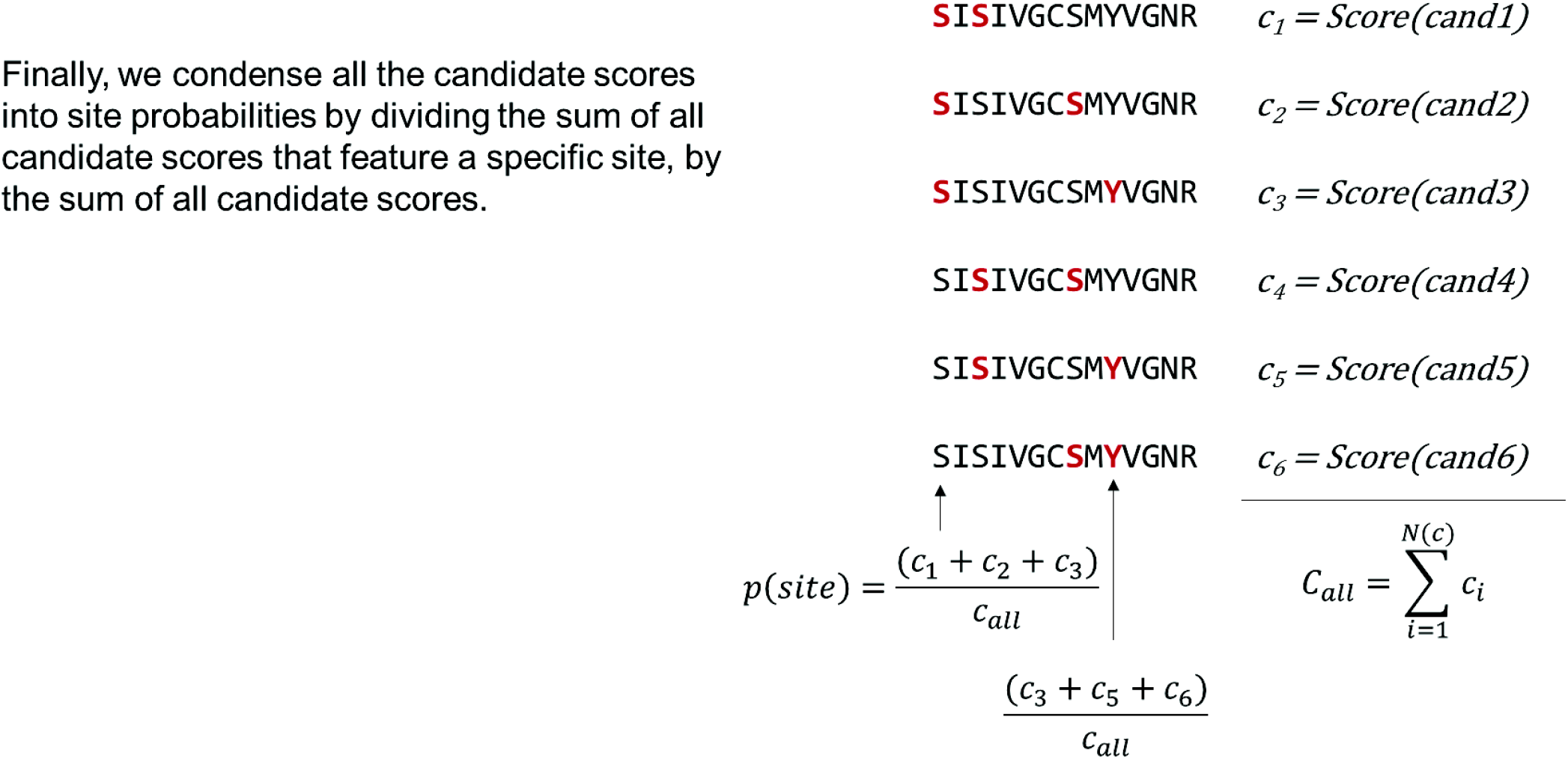

**Supplementary Note 1:**
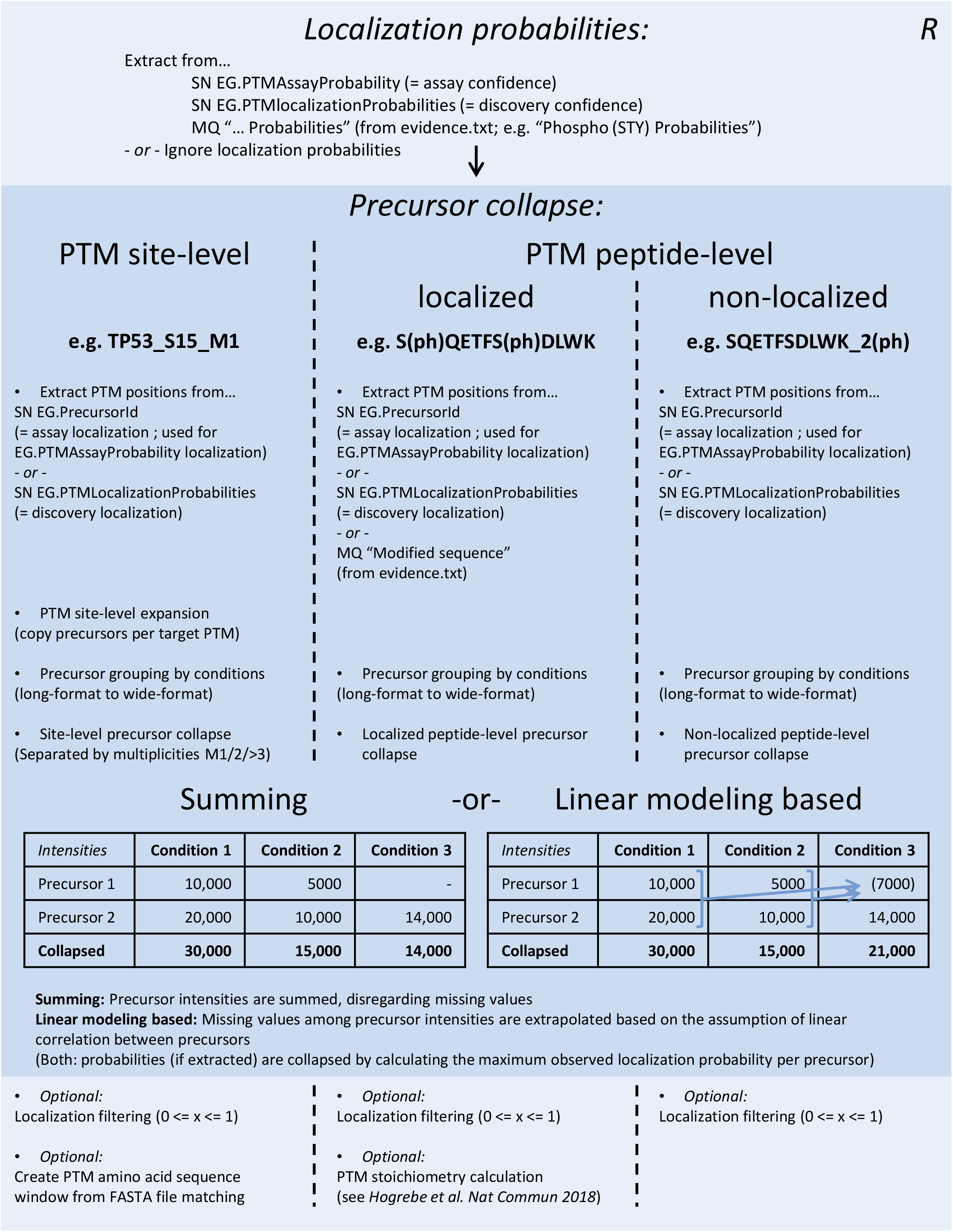
Perseus plugin peptide collapse. This is a schematic of the data processing steps of the Perseus plugin “Peptide Collapse”. The main purpose of the plugin is to combine precursor quantifications into consensus PTM sites or peptides. This should allow more robust statistical analysis, especially with missing values present in the data. Precursor collapse levels mimic the MaxQuant site, evidence (= Modified sequence) and modification-specific-peptides output tables.

## REFERENCES (Up to 40)

1. Olsen, J. V. & Mann, M. Status of large-scale analysis of post-translational modifications by mass spectrometry. Mol. Cell. Proteomics 12, 3444–3452 (2013).

2. Olsen, J. V. et al. Global, in vivo, and site-specific phosphorylation dynamics in signaling networks. Cell 127, 635–648 (2006).

3. Villén, J., Beausoleil, S. A., Gerber, S. A. & Gygi, S. P. Large-scale phosphorylation analysis of mouse liver. Proc. Natl. Acad. Sci. U. S. A. 104, 1488–1493 (2007).

4. Lundby, A. et al. In vivo phosphoproteomics analysis reveals the cardiac targets of β-adrenergic receptor signaling. Sci. Signal. 6, rs11 (2013).

5. Francavilla, C. et al. Multilayered proteomics reveals molecular switches dictating ligand-dependent EGFR trafficking. Nat. Struct. Mol. Biol. 23, 608–618 (2016).

6. Röst, H. L. et al. OpenSWATH enables automated, targeted analysis of data-independent acquisition MS data. Nat. Biotechnol. 32, 219–223 (2014).

7. Bruderer, R. et al. Extending the limits of quantitative proteome profiling with data-independent acquisition and application to acetaminophen-treated three-dimensional liver microtissues. Mol. Cell. Proteomics 14, 1400–1410 (2015).

8. Kelstrup, C. D. et al. Performance Evaluation of the Q Exactive HF-X for Shotgun Proteomics. J. Proteome Res. 17, 727–738 (2018).

9. Oda, K., Matsuoka, Y., Funahashi, A. & Kitano, H. A comprehensive pathway map of epidermal growth factor receptor signaling. Mol. Syst. Biol. 1, 2005.0010 (2005).

10. Olsen, J. V. et al. Higher-energy C-trap dissociation for peptide modification analysis. Nat. Methods 4, 709–712 (2007).

11. Parker, B. L. et al. Targeted phosphoproteomics of insulin signaling using data-independent acquisition mass spectrometry. Sci. Signal. 8, rs6 (2015).

12. Hoffman, N. J. et al. Global Phosphoproteomic Analysis of Human Skeletal Muscle Reveals a Network of Exercise-Regulated Kinases and AMPK Substrates. Cell Metab. 22, 922–935 (2015).

13. Peckner, R. et al. Specter: linear deconvolution for targeted analysis of data-independent acquisition mass spectrometry proteomics. Nat. Methods (2018). doi:10.1038/nmeth.4643

14. Rosenberger, G. et al. Inference and quantification of peptidoforms in large sample cohorts by SWATH-MS. Nat. Biotechnol. 35, 781–788 (2017).

15. Meyer, J. G. et al. PIQED: automated identification and quantification of protein modifications from DIA-MS data. Nat. Methods 14, 646–647 (2017).

16. Tusher, V. G., Tibshirani, R. & Chu, G. Significance analysis of microarrays applied to the ionizing radiation response. Proc. Natl. Acad. Sci. U. S. A. 98, 5116–5121 (2001).

17. Cox, J. et al. Andromeda: a peptide search engine integrated into the MaxQuant environment. J. Proteome Res. 10, 1794–1805 (2011).

18. Cox, J. & Mann, M. MaxQuant enables high peptide identification rates, individualized p.p.b.-range mass accuracies and proteome-wide protein quantification. Nat. Biotechnol. 26, 1367–1372 (2008).

19. Batth, T. S., Francavilla, C. & Olsen, J. V. Off-line high-pH reversed-phase fractionation for in-depth phosphoproteomics. J. Proteome Res. 13, 6176–6186 (2014).

20. Sharma, K. et al. Ultradeep human phosphoproteome reveals a distinct regulatory nature of Tyr and Ser/Thr-based signaling. Cell Rep. 8, 1583–1594 (2014).

21. Bekker-Jensen, D. B. et al. An Optimized Shotgun Strategy for the Rapid Generation of Comprehensive Human Proteomes. Cell Syst 4, 587–599.e4 (2017).

22. Olsen, J. V. et al. Quantitative phosphoproteomics reveals widespread full phosphorylation site occupancy during mitosis. Sci. Signal. 3, ra3 (2010).

23. Presler, M. et al. Proteomics of phosphorylation and protein dynamics during fertilization and meiotic exit in the Xenopus egg. Proc. Natl. Acad. Sci. U. S. A. 114, E10838–E10847 (2017).

24. Hogrebe, A. et al. Benchmarking common quantification strategies for large-scale phosphoproteomics. Nat. Commun. 9, 1045 (2018).

25. Fabregat, A. et al. The Reactome Pathway Knowledgebase. Nucleic Acids Res. 46, D649–D655 (2018).

26. Post, H. et al. Robust, Sensitive, and Automated Phosphopeptide Enrichment Optimized for Low Sample Amounts Applied to Primary Hippocampal Neurons. J. Proteome Res. 16, 728–737 (2017).

27. Gessulat, S. et al. Prosit: proteome-wide prediction of peptide tandem mass spectra by deep learning. Nat. Methods (2019). doi:10.1038/s41592-019-0426-7

28. Tiwary, S. et al. High-quality MS/MS spectrum prediction for data-dependent and data-independent acquisition data analysis. Nat. Methods (2019). doi:10.1038/s41592-019-0427-6

29. Meier, F. et al. Online parallel accumulation – serial fragmentation (PASEF) with a novel trapped ion mobility mass spectrometer. Mol. Cell. Proteomics mcp.TIR118.000900 (2018).

30. Rappsilber, J., Mann, M. & Ishihama, Y. Protocol for micro-purification, enrichment, pre-fractionation and storage of peptides for proteomics using StageTips. Nat. Protoc. 2, 1896–1906 (2007).

